# Structure and mechanism of a cytosine transport protein

**DOI:** 10.1101/2021.12.23.473980

**Authors:** Caitlin E. Hatton, Deborah H. Brotherton, Alexander D. Cameron

## Abstract

CodB is a cytosine transporter from the Nucleobase-Cation-Symport-1 (NCS1) transporter family, a member of the widespread LeuT superfamily. Previous experiments with the nosocomial pathogen *Pseudomonas aeruginosa* have shown CodB also to be important in the uptake of 5-fluorocytosine, which has been suggested as a novel drug to combat antimicrobial resistance by suppressing virulence in the organism. Here we solve the crystal structure of CodB from *Proteus vulgaris*, at 2.4Å resolution in complex with cytosine. We show that the protein carries out the sodium-dependent uptake of cytosine and can bind 5-fluorocytosine. Comparison of the substrate-bound structures of CodB and the hydantoin transporter Mhp1, the only other NCS1 family member for which the structure is known, highlight the importance of the hydrogen bonds that the substrates make with the main chain at the breakpoint in the discontinuous helix, TM6. In contrast to other LeuT superfamily members, neither CodB nor Mhp1 make specific interactions with residues on TM1. Comparison of the structures provides insight into the intricate mechanisms of how these proteins transport substrates across the plasma membrane.

## Introduction

The cytosine transporter CodB belongs to the nucleobase cation symporter 1 (NCS1) family of membrane transporters (de Koning & Diallinas, 2000). The NCS1 family is found in bacteria (de Koning & Diallinas, 2000), archaea (Ma *et al*, 2013), fungi (Pantazopoulou & Diallinas, 2007) and plants (Mourad *et al*, 2012; Schein *et al*, 2013; Witz *et al*, 2014). Members of the family are responsible for transporting nucleobases and related molecules into cells, often as components of salvage pathways. CodB is found in an operon with CodA, a cytosine deaminase, which converts cytosine to uracil and ammonia, providing an alternative nitrogen source (Fig 1) (Danielsen *et al*, 1992). In the nosocomial pathogen *Pseudomonas aeruginosa*, CodB has been shown to be important in the effect of 5-fluorocytosine in the suppression of virulence (Imperi *et al*, 2013). 5-fluorocytosine is initially taken up by CodB and then converted to toxic 5-fluorouracil by CodA, which in turn represses the production of bacterial virulence factors resulting in reduced pathogenicity in mouse models of infection (Imperi *et al*., 2013). 5-fluorocytosine is already used in the clinic as an antimycotic drug (Vermes *et al*, 2000), the toxicity of 5-fluorouracil being avoided because cytosine deaminases are not found in higher eukaryotes. Drugs that cause a reduction in virulence rather than growth may present a novel means to combat antibiotic resistance as they may not exert the same selective pressure on the organism to develop resistance as traditional antibiotics (Reviewed by (Maura *et al*, 2016)).

**Fig 1:**
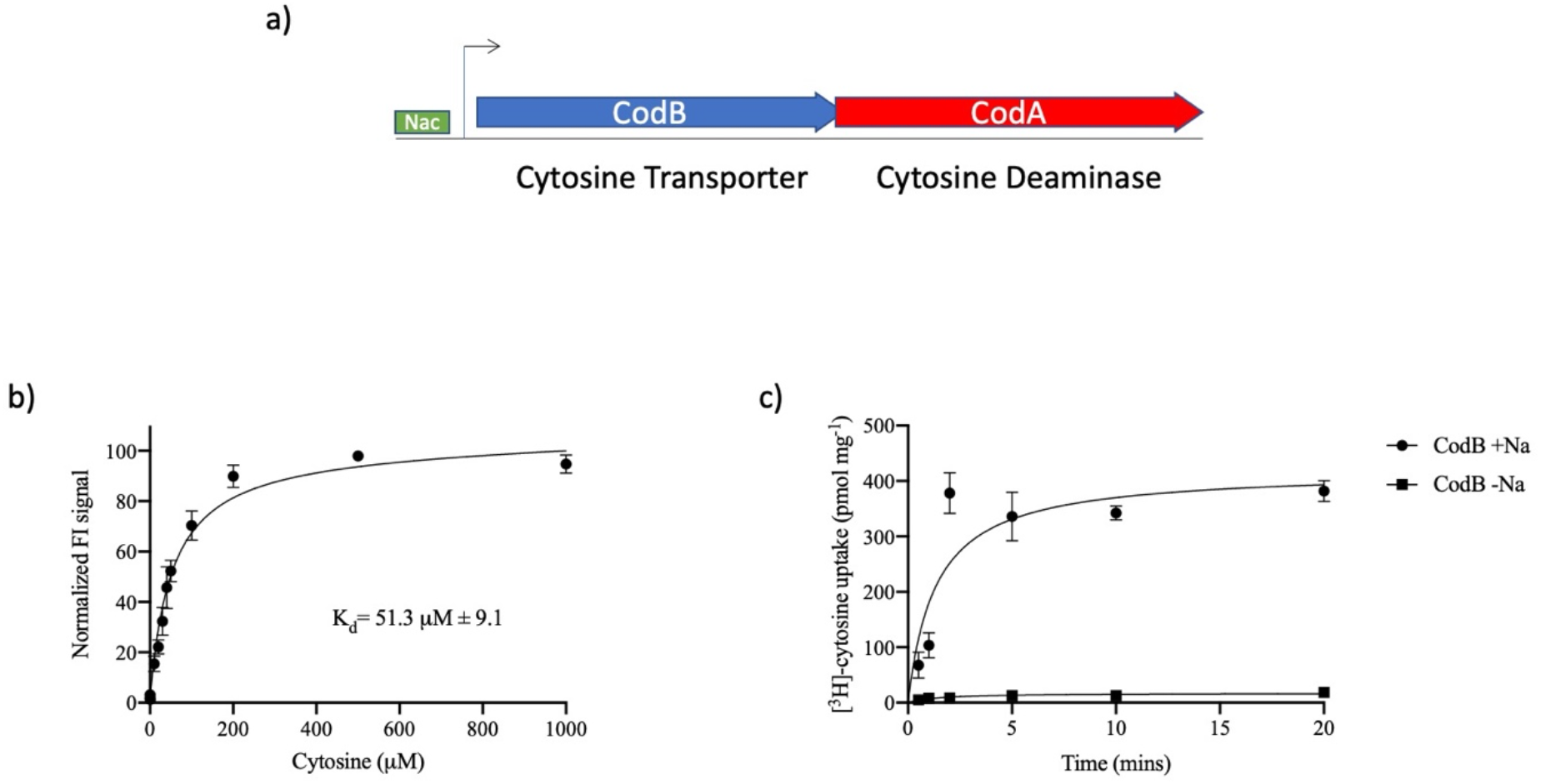
Cytosine binding and transport by CodB. **a)** CodB is found in an operon with CodA with overlapping genes. Transcription is regulated by the Nitrogen Assimilation Control protein (NAC) in response to low nitrogen levels (Danielsen *et al*., 1992; Muse *et al*, 2003; Santos-Zavaleta *et al*, 2019). **b)** Binding affinity of CodB for cytosine as measured using the thermostability assay. Cytosine was titrated into detergent solubilised membranes from cells overexpressing CodB. The K_d_ was estimated to be 51 ± 9 µM. The measurements are the average of 4 independent titrations with error bars of the s.e.m. **b)** Time course of ^3^H-5-cytosine uptake by CodB. Experiments were done, either in the presence or absence of an inwardly-directed sodium ion gradient. Lemo21 (DE3) cells were used as a background measurement, with Lemo21(DE3) expressing CodB. Values reported are the averaged mean ± s.e.m. from n = 3 independent cultures.

CodB has 24% sequence identity to the sodium-dependent hydantoin transporter, Mhp1 from *Mycobacterium liquefaciens*, the only member of the NCS1 family for which the structure is known (Kazmier *et al*, 2014a; Shimamura *et al*, 2010; Simmons *et al*, 2014; Weyand *et al*, 2008). Determination of the structure of Mhp1 placed the NCS1 family in the amino acid polyamine organocation (APC) transporter or LeuT superfamily (Wong *et al*, 2012). Mhp1, like other members of this superfamily has a common core built of a pseudosymmetric 5 transmembrane helix, inverted repeat (Abramson & Wright, 2009) with the two repeating units intertwining to give two domains. The first of these domains, the bundle consists of TM1-2 and TM6-7 and is characterised by two discontinuous helices (TM1 and TM6). Amongst the LeuT superfamily the second domain is commonly referred to as the scaffold domain, (TM3-5, TM8-10) (Forrest *et al*, 2008), however, in Mhp1 TM5 and TM10 appear to act partially independently of the scaffold domain and so the reduced domain (TM2-4 and TM8-9) has been called the hash domain. Substrates for the respective transporters bind at the interface of the bundle and hash domains near to the breakpoints of the two discontinuous helices of the bundle domain. Secondary transporters work by the alternating access mechanism in which the binding site of the protein alternatively faces one side of the membrane or the other (Jardetzky, 1966). The structure of Mhp1 has been solved in the three main states associated with alternating access: outward-facing with sodium bound (Weyand *et al*., 2008); outward-facing occluded with sodium and substrate bound (Simmons *et al*., 2014; Weyand *et al*., 2008) and inward-facing (Shimamura *et al*, 2008). In transitioning between the outward-facing and inward-facing states the hash domain moves relative to the bundle domain as an approximate rigid body (Kazmier *et al*., 2014a; Shimamura *et al*., 2010). This mechanism, which is supported by studies using DEER (Kazmier *et al*., 2014a), largely conforms to the rocking bundle model that was first proposed for the leucine transporter LeuT, the founding member of the LeuT superfamily, based on an analysis of the symmetry repeats of the protein (Forrest *et al*., 2008). The occlusion of the substrate in the binding pocket from the extracellular side prior to transport and the inferred opening of the site to the inward-side is effected by the flexing of two symmetry related helices, TM10 and TM5 respectively (Malinauskaite *et al*, 2014; Shimamura *et al*., 2010; Weyand *et al*., 2008), with a short extracellular helix (EL4) sealing the site as it transitions to the inward-facing side (Kazmier *et al*., 2014a; Shimamura *et al*., 2010).

Of the members of the LeuT superfamily that have been solved to date, several, like Mhp1 are sodium coupled. These include LeuT (Yamashita *et al*, 2005), MhsT (Malinauskaite *et al*., 2014), dDAT (Penmatsa *et al*, 2013), SERT (Coleman *et al*, 2016) and GlyT (Shahsavar *et al*, 2021) of the neurotransmitter sodium symporters (NSS) family, vSGLT (Faham *et al*, 2008), SGLT (Han *et al*, 2021; Niu *et al*, 2021) and SiaT (Wahlgren *et al*, 2018) from the solute sodium symporters (SSS) and BetP (Ressl *et al*, 2009) from the betaine/choline/carnitine transporters (BCCTs) family. While the stoichiometry of sodium ions varies amongst the different proteins, the sodium site that is observed in Mhp1 is conserved in all. This site (known as Na2 following its nomenclature in the structure of LeuT) is coordinated by residues at the breakpoint of TM1 of the bundle domain and residues on TM8 of the hash motif. Intuitively, therefore, the conserved sodium site is located at a position that is ideal for stabilising the outward-facing state of the protein. With respect to these other transporters, Mhp1 is unusual in two respects. Firstly, whereas in the other proteins the respective substrates make critical interactions with the breakpoint of TM1, in Mhp1 there are no direct hydrogen bonding interactions between the substrate and TM1, at least as modelled at the limited resolution (3.4Å) of the substrate-bound structures. Secondly, whereas studies of the other superfamily members show the position of TM1 varies dependent on the conformational state of the protein (Coleman *et al*, 2019; Kazmier *et al*, 2014b; Krishnamurthy & Gouaux, 2012; Perez *et al*, 2012), in Mhp1 these movements are much more subtle (Kazmier *et al*., 2014a; Shimamura *et al*., 2010; Simmons *et al*., 2014).

CodB transports cytosine, a much smaller compound than the bulky substituted hydantoins transported by Mhp1. To understand how cytosine and 5-fluorocytosine bind in the substrate binding site we solve the crystal structure of the protein in complex with cytosine and a sodium ion at 2.4Å resolution. Combining this data with transport assays and site-directed mutagenesis provides insight into molecular recognition and transport in CodB and the NCS1 family and indeed the APC superfamily in general.

## Results

### CodB is a Sodium-Dependent Cytosine Transporter

CodB from the opportunistic pathogen *Proteus vulgaris* (CodB_PV_) was identified as suitable for structural studies using fluorescent-based screening methods (Drew *et al*, 2006; Sonoda *et al*, 2011). CodB_PV_ has 84% sequence identity with CodB from *Escherichia coli* and 74% identity with that from *P aeruginosa* (Supplementary Figure 1). In a stabilisation assay (Nji *et al*, 2018) cytosine was observed to stabilise the detergent solubilised protein (Fig 1a). Using stabilisation as a surrogate for binding, the affinity was measured to be ∼50 μM. Although there are no reports of sodium-dependency in CodB, given that the residues involved in sodium ion coordination are conserved between CodB and Mhp1 (Supplementary Figure 1) we suspected that, like Mhp1, the transporter would be sodium coupled. Sodium-dependent uptake of cytosine was confirmed using an in-cell transport assay by following the uptake of ^3^H-cytosine (Fig 1b).

### Overall Structure and Conformation of CodB

CodB was purified and crystallised in the presence of cytosine using the lipidic cubic phase (LCP) method (Caffrey & Cherezov, 2009) and the structure determined and refined at 2.4Å to an Rfactor of 20.1% and a corresponding Rfree of 24.6% (Table 1). The addition of cytosine during purification was observed to reduce protein loss, consistent with stabilisation of the protein. CodB crystallises as a monomer with two molecules in the asymmetric unit orientated oppositely with respect to the membrane plane. Both molecules adopt an outward-open conformation (Fig 2, Supplementary Figure 2) with cytosine bound in a solvent accessible polar pocket and density consistent with a Na^+^ ion observed in the conserved Na2 site. The overall 12-TM helix topology is very similar to Mhp1: TM1-TM5 are related to TM6-TM10 by a pseudo 2-fold axis and intertwine to form the bundle and hash motifs (Fig 2c). Transmembrane helices TM11-TM12 abut the hash motif. Between EL4 and the tip of TM3, TM10 and TM1 there is non-protein density, which we have tentatively modelled as DDM and monoolein respectively (Supplementary Fig 2). Overall, the root mean square deviation (RMSD) between CodB and the occluded form of Mhp1 is 2Å for 344 Cα atoms out of a possible 416 (Fig 3). TM8 forms a more regular helix than seen in Mhp1 where there is a single residue insertion into the helix next to the substrate binding site (Fig 3b, Supplementary Figure 1), but most of the substantial differences are in the loop regions where the sequence of CodB is generally shorter than Mhp1 (Fig 3, Supplementary Figure 1). The helix that forms part of EL4, which is critical in sealing the extracellular cavity on the transition to the inward-facing form (Kazmier *et al*., 2014a; Shimamura *et al*., 2010) in Mhp1 is set more deeply into the cavity (Fig 3a). In Mhp1 TM10 bends towards the substrate upon substrate binding. In CodB this TM is in a position more reminiscent of the non-substrate bound outward-open form of Mhp1 rather than the substrate occluded form (Fig 3b and c).

**Table 1:**
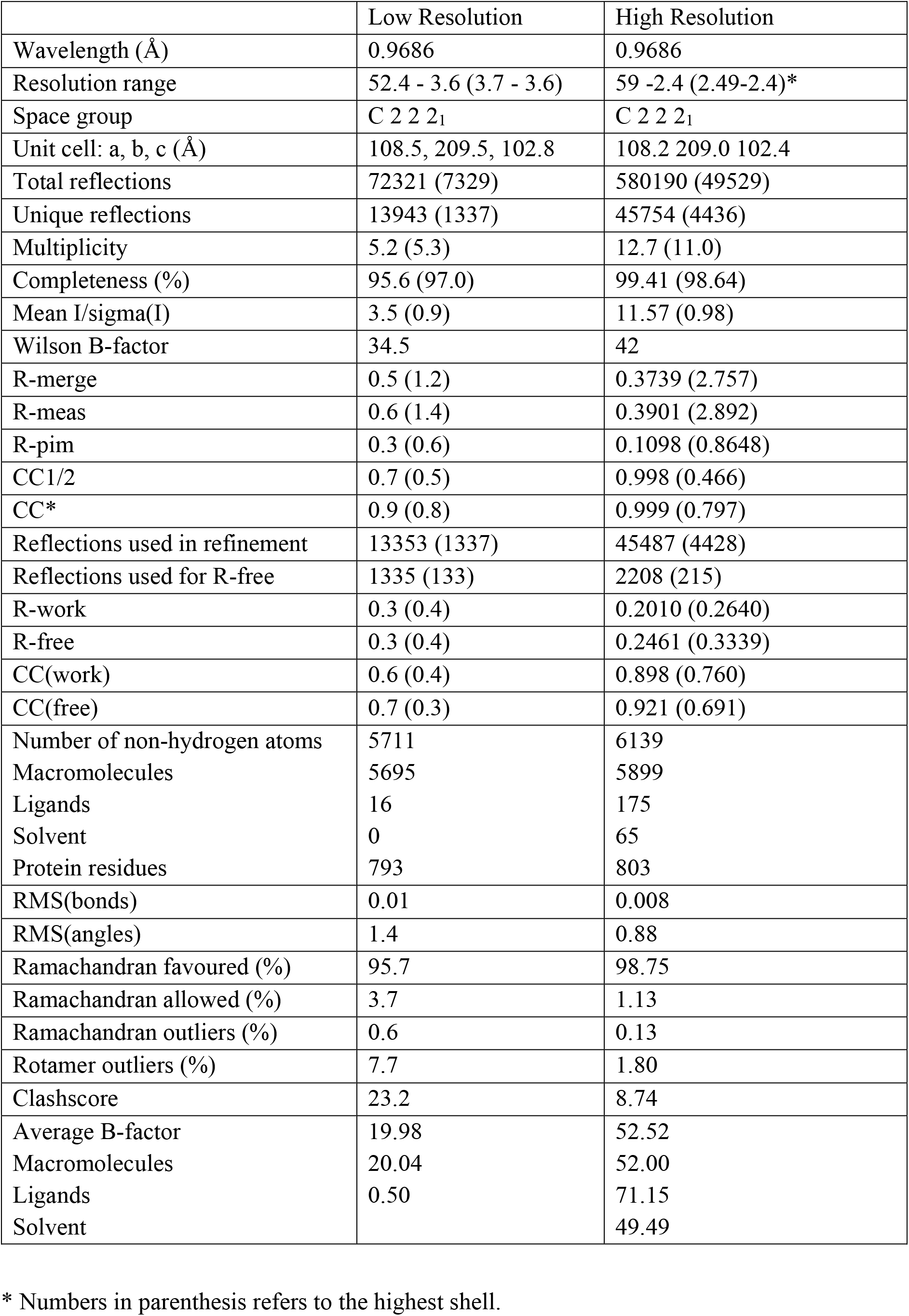
Data Processing and Refinement Statistics.

**Fig 2:**
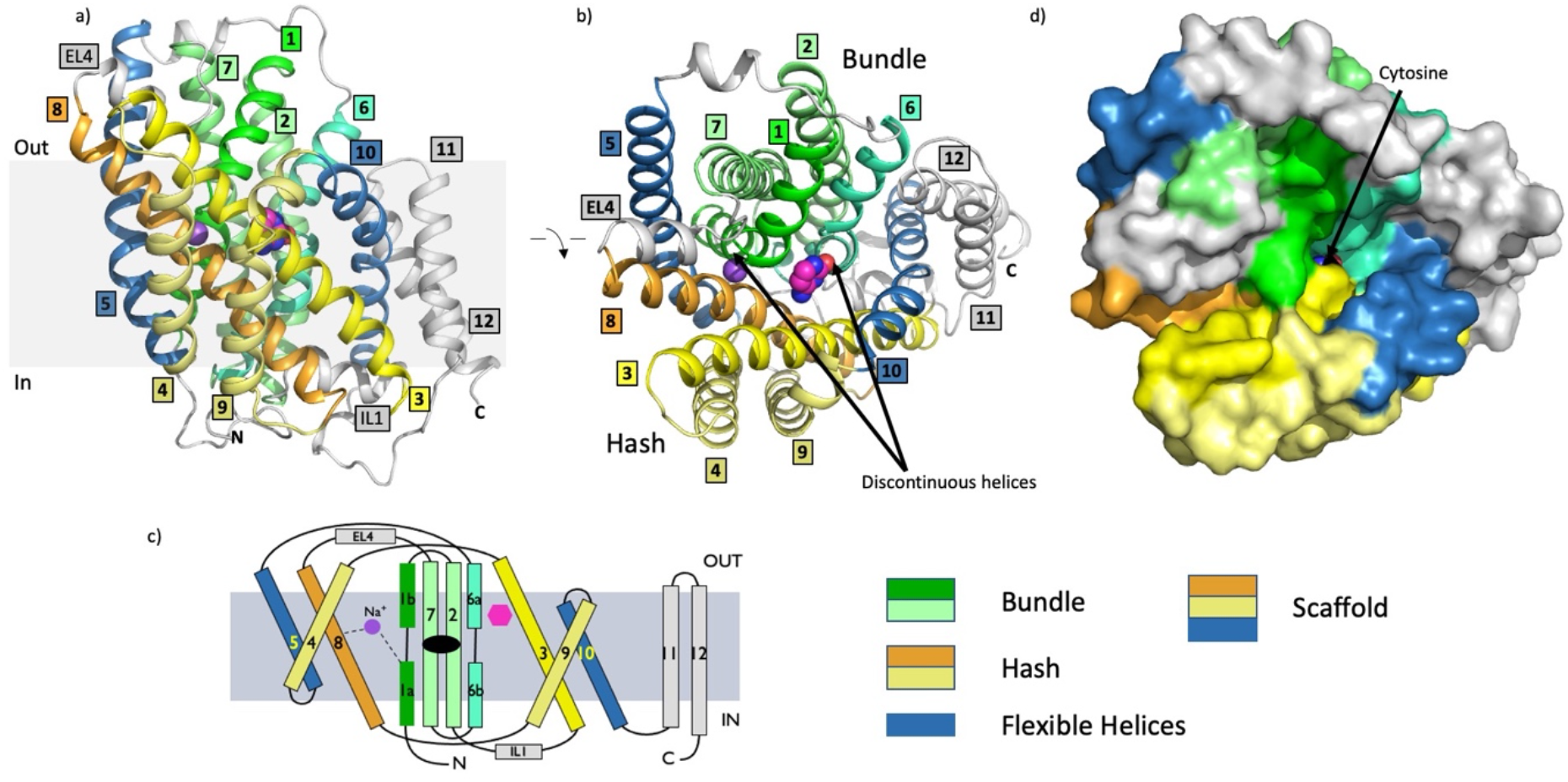
Structure of CodB. **a)** Ribbon diagram of CodB in the plane of the membrane. The bundle motif is depicted in different shades of green with TM1 in green, TM6 in sea green and TMs 2 and 7 in light green. The hash motif is shown with TM3 in yellow, TM8 in orange and TMs 4 and 9 in light yellow. The flexible helices TM5 and TM10 have been coloured blue. In other LeuT superfamily members the combination of the hash motif and the flexible helices are often referred to as the scaffold domain (Forrest *et al*., 2008). TM11 and TM12 are coloured grey. EL4 is the extracellular loop linking TMs 7 and 8 and IL1 is the intracellular loop between TMs 2 and 3. The carbon atoms of the cytosine are coloured magenta. The sodium ion is depicted as a purple sphere. **b)** As (a) but looking from the extracellular side of the membrane. **c)** Topology diagram coloured as in a. **d)** Surface representation in the same colouring as a and b with the same view as (b). The cytosine can be observed at the bottom of an open cavity.

**Fig 3:**
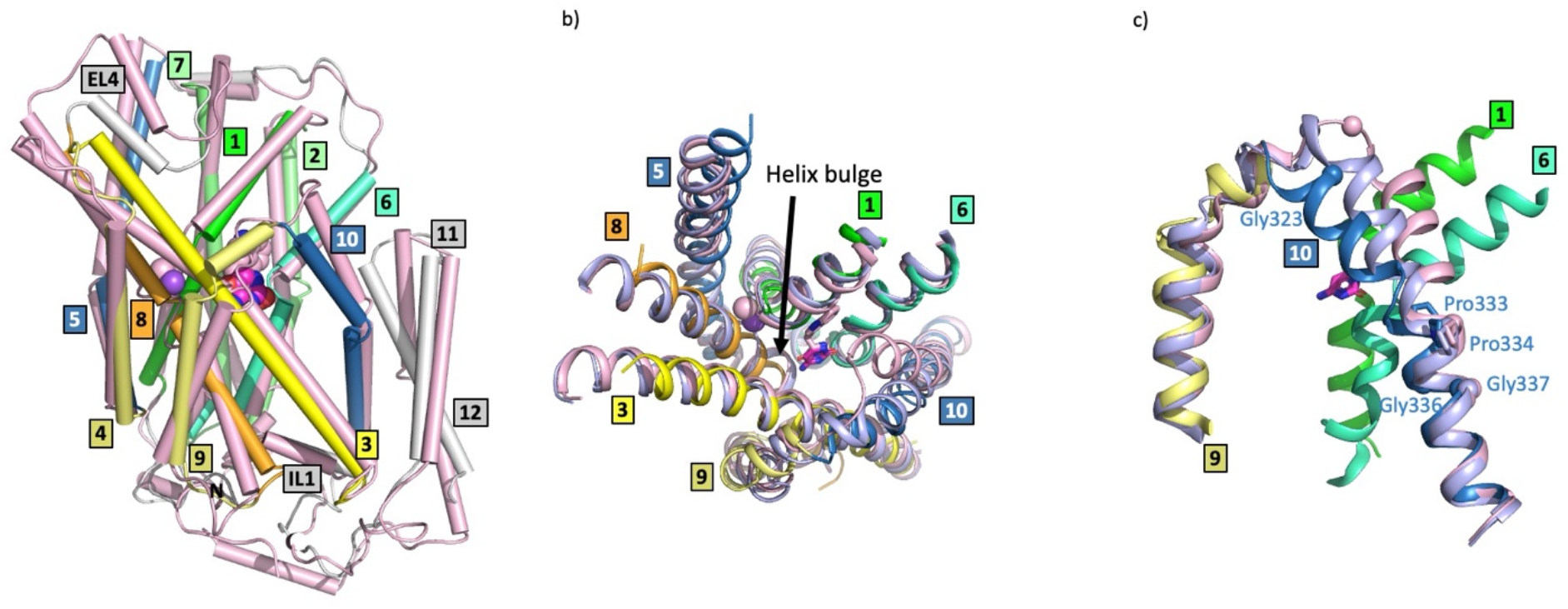
Comparison of the structures of CodB and Mhp1. **a)** Overlay of the complete structures with CodB coloured as in Figure 2 and the occluded form of Mhp1 in pink. Comparing CodB with the outward-occluded form of Mhp1 (4D1A), CodB aligned with a RMSD of 2.0Å for 344 residues out of a possible 416. These differences are distributed throughout the protein. When the 4-helix bundle and hash domain were extracted and aligned the respective RMSDs were 2.1Å for 123 out of 152 residues for the bundle domain and 1.7Å for 99 out of 119 residues for the hash domain. **b)** View of the sodium and cytosine/hydantoin binding sites from the extracellular side. **c)** Comparison of TM10 in the structures of CodB (cyan), the outward-open form of Mhp1 (2JLN lilac) and the occluded form of Mhp1 (4D1A pink). TMs 9 and 10 are shown for all structures. For CodB TMs 1 and 6 are also shown to help orientation. Glycines are denoted by spheres at the C_α_ atoms and prolines as sticks. The glycine and proline residues on TM10 of CodB are labelled.

### Cytosine Binding Site

The cytosine substrate is found at the interface of the hash-motif and the 4-helix bundle, sandwiched between two aromatic residues, Trp108 of TM3 of the hash domain and Phe204 of TM6 of the bundle domain (Fig 4) in a face-to-face pi stacking arrangement. The cytosine forms two direct hydrogen bonds to the mainchain at either side of the breakpoint of TM6 of the bundle domain: to the carbonyl oxygen of Ser203 of TM6a and to the main chain nitrogen atom of Ala207 of TM6b (Fig 4a). In addition, a water molecule bridges the cytosine to the carbonyl oxygens of Gly202 (TM6a) and Ser 206 (TM6b). In terms of interactions with the hash motif, as well as stacking with Trp108, there is a hydrogen bond from the cytosine to the amino oxygen of Gln105 of TM3 and a potential water-mediated hydrogen bond to Asn280 of TM3.

**Fig 4:**
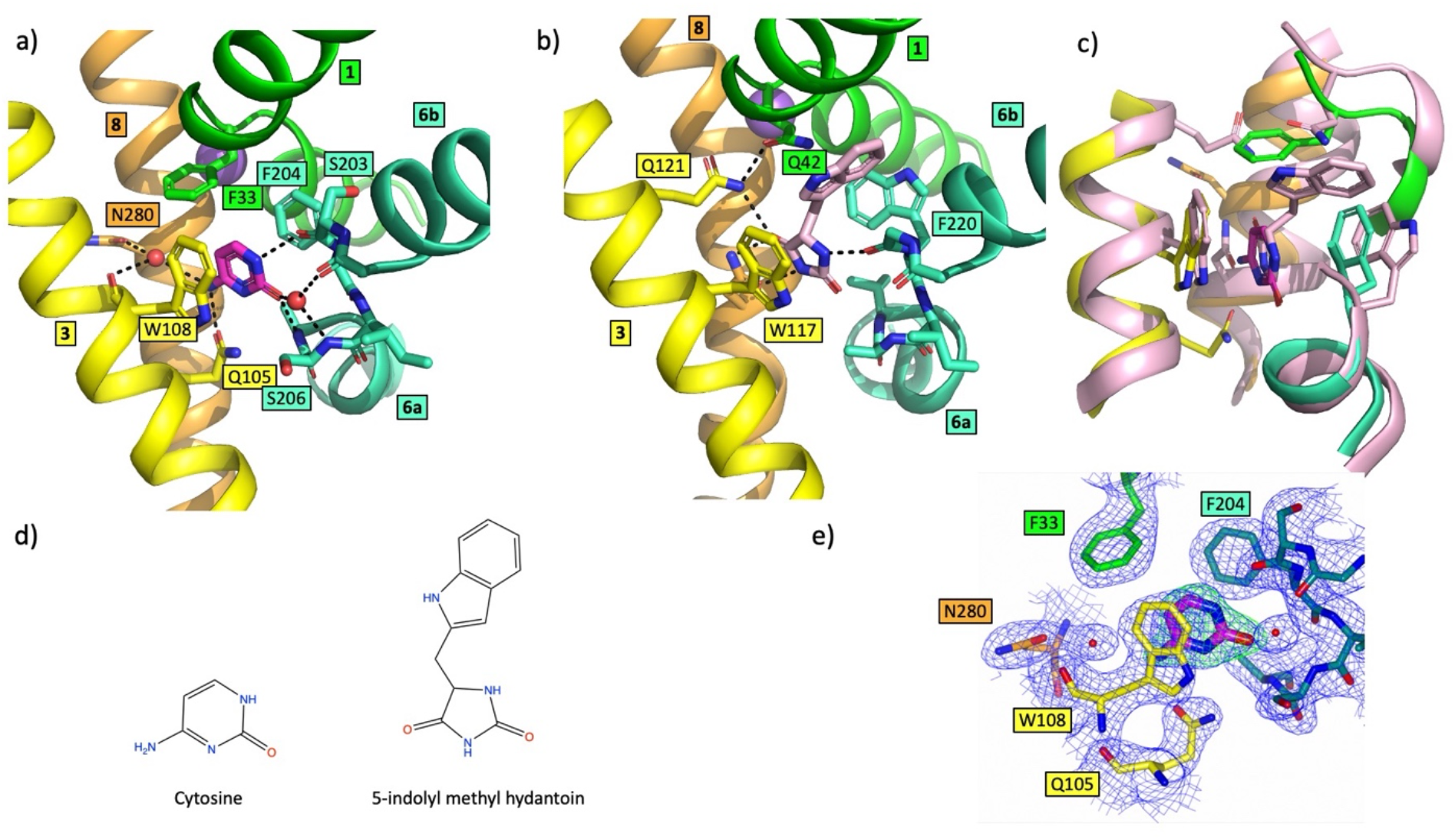
Comparison of the substrate binding sites in CodB and Mhp1. **a)** View of the binding site of CodB coloured as in Figure 1. Water molecules are denoted as red spheres and potential hydrogen bonds are shown as dashed lines. **b)** Mhp1 in the same view as (a) and coloured as for CodB. The indolylmethyl-hydantoin is shown with pink carbon atoms. **c)** Overlay of CodB and Mhp1 with CodB coloured as in (a) and Mhp1 in pink. **d)** Chemical structures of the respective substrateDFc map is based on phases from the refineds **e)** Electron density for the cytosine. The 2mFo-DFc map in blue is contoured at 1σ and the mFo-DFc map, calculated before the addition of the cytosine, in green at 3σ.

When the substrate binding site in CodB is compared to Mhp1 and other members of the NCS1 family the relative importance of the interactions that the base makes with the protein can be inferred. The two aromatic residues, which sandwich the cytosine in CodB are conserved throughout the NCS1 family; whereas the equivalent of Trp108 is predominantly a tryptophan, the equivalent of Phe204 can be either a phenylalanine as seen in CodB or a tryptophan as in Mhp1. The remaining residues that interact with the cytosine in CodB are much less conserved throughout the family. In Mhp1 the major hydrogen bonding interactions with the substrate involving the residues from the hash motif are with Asn318 from TM8 and Gln121 from TM3 rather than the equivalent of Gln105. Both substrates, from CodB and Mhp1, however, are within hydrogen bonding distance of TM6. Whereas in CodB the cytosine interacts with the main chain atoms of both TM6a and TM6b on either side of the helix break, in Mhp1 only TM6a is within hydrogen bonding distance of the hydantoin. The equivalent interaction between the substrate and TM6b to that seen in CodB is ∼3.8Å, slightly too long for a hydrogen bond, although it is possible that this also reflects the resolution of the Mhp1 structure (3.4Å) and with minor adjustments of the positioning of the base and/or the main chain atoms could bring the two atoms to a position more consistent with a hydrogen bond. What is remarkable is that when the structure of CodB is superposed on that of Mhp1 based on their respective Cα atoms, the cytosine of CodB and the hydantoin moiety of the Mhp1 substrate overlap almost exactly (Fig 4). This is surprising given that firstly, the substrates of the two proteins are different, and secondly, there is limited conservation within the binding sites. The fact that the cytosine and the hydantoin moiety of the respective substrates overlap so well demonstrates the importance of the interactions with TM6 as well as the aromatic residues. It is noteworthy that after superposing the two proteins as above, the phenyl ring of CodB overlaps the 6-membered ring of the tryptophan in Mhp1 (Fig 4).

Neither CodB nor Mhp1 have specific interactions involving the respective substrates and TM1. The only interaction that the cytosine makes with TM1 is a potential edge to face pi-stacking arrangement with Phe33. The equivalent residue in Mhp1 is Gln42, which is not involved in hydrogen bonding the substrate, but is within hydrogen bonding distance of Gln121.

### Sodium Binding Site

Electron density consistent with a sodium ion is visible at the Na2 site that is conserved amongst the Na^+^-coupled LeuT transporters (Fig 5). The sodium ion is coordinated by the main chain carbonyl oxygens of Gly29 and Phe32 at the breakpoint of TM1 of the bundle domain and the main chain carbonyl oxygen of Asn275 and the hydroxyl oxygens of Thr278 and Thr279 from TM8 of the hash domain in a square pyramidal arrangement (Fig 5). In an unusual interaction that is not seen in other Na^+^-coupled LeuT members, the side chain of Asn275 is also within hydrogen-bond distance of the side chain hydroxyl and the amide nitrogen of Ser34 at the C-terminus of TM1b, providing a further link between the hash and bundle domains when sodium binds (Fig 5). More typically hydrophobic residues are found at this position. Interestingly, Asn282, also on TM8 and positioned just below the sodium ion, in the view shown in Fig 5, also forms hydrogen bonding interactions to Val26 of TM1a bridging these two helices (Fig 5).

**Fig 5:**
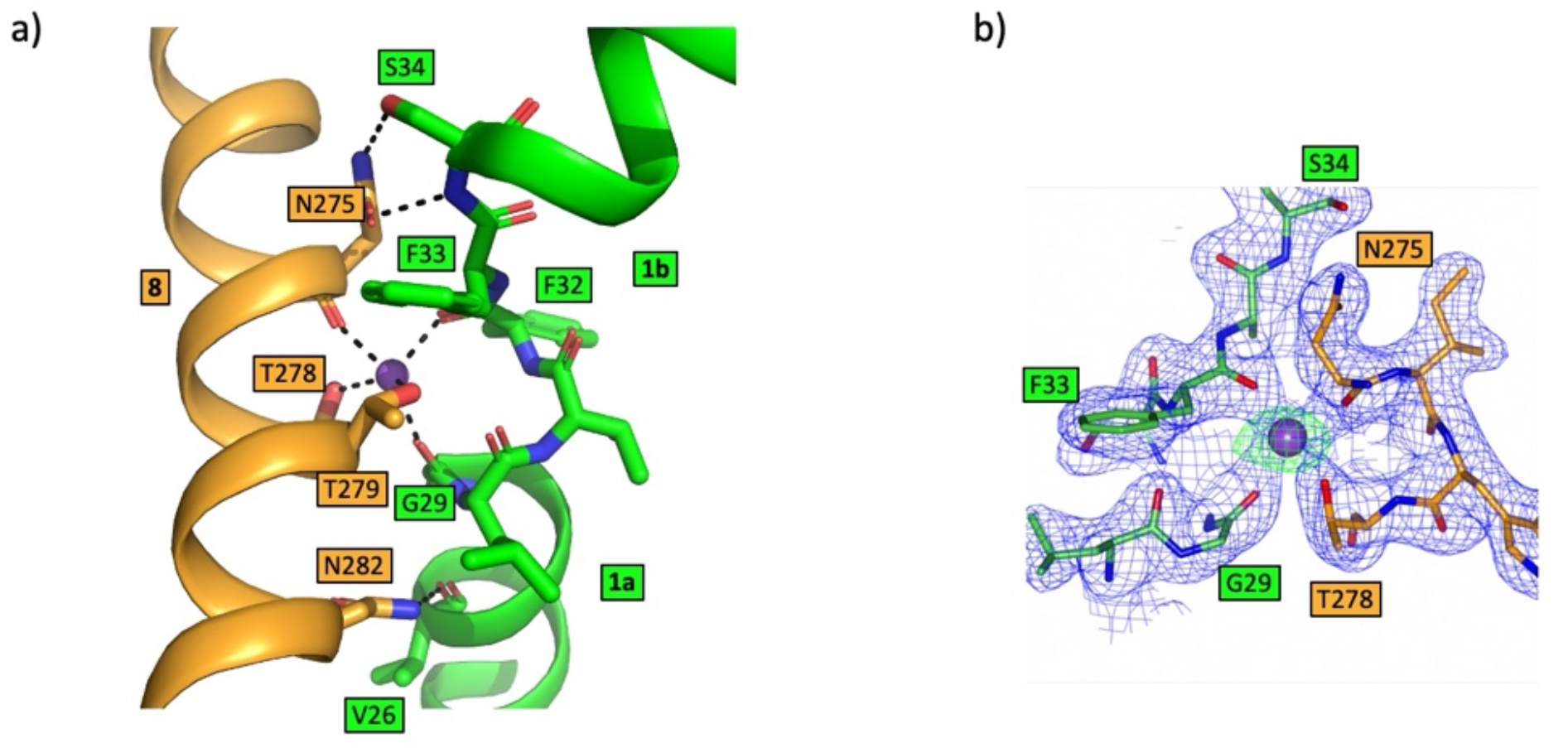
Sodium binding site in CodB. **a)** View showing the interactions between TM1 and TM8 centred on the sodium ion. The sodium ion makes interactions with residues on TM1 and TM8 (black dashed lines). Asn275 and Asn282 are also within hydrogen bonding distance of residues on TM1. **b)** Electron density associated with the sodium ion. The 2mFo-DFc map in blue is contoured at 1σ and the mFo-DFc map, calculated before the addition of the sodium ion is in green at 5σ.

### Molecular recognition in CodB

The three key residues that interact with the cytosine through their side chains are Gln105, Trp108 and Phe204. While mutation of any of these residues to alanine caused an apparent reduction in binding of the cytosine, as monitored through the stabilisation assay (Supplementary Fig 3), only mutations of Gln105 or Trp108 caused a dramatic reduction in the transport of ^3^H-cytosine (Fig 6). Mutation of Asn280, which forms a water-mediated hydrogen bond had no effect on transport, though it did appear to affect binding (Fig 6 and Supplementary Figure 3). To investigate the specificity of CodB for cytosine a selection of nucleobases and related compounds were tested for their effect on stabilising the protein or in inhibiting transport of ^3^H-cytosine. Of the bases investigated, cytosine was the most effective both at stabilising the protein and inhibiting transport (Fig 6 and Supplementary Figure 4). Consistent with its effect on *P. aeruginosa* (Imperi *et al*., 2013) 5-fluorocytosine also showed some inhibition of cytosine uptake (Fig 6) with a K_D_ estimated from the stability assay of 285μM (Supplementary Figure 4). It would seem likely that this would bind in a similar mode to cytosine with the fluorine interacting with Phe33. Methylcytosine, where the fluorine is replaced with a much larger methyl group on the other hand doesn’t bind, presumably because the methyl group is likely to clash with Phe33. Although, both uracil and isocytosine inhibited the uptake of ^3^H-cytosine under the conditions of the uptake assay, they had little effect in stabilising the protein in the stabilisation assay (Supplementary Fig 4). Modelling of the uracil into the pocket, based on the cytosine binding mode, suggests that the uracil may be able to bind if the side chain of Gln105 were to flip and this may also occur with isocytosine. No binding was observed for the pyrimidine bases.

**Fig 6:**
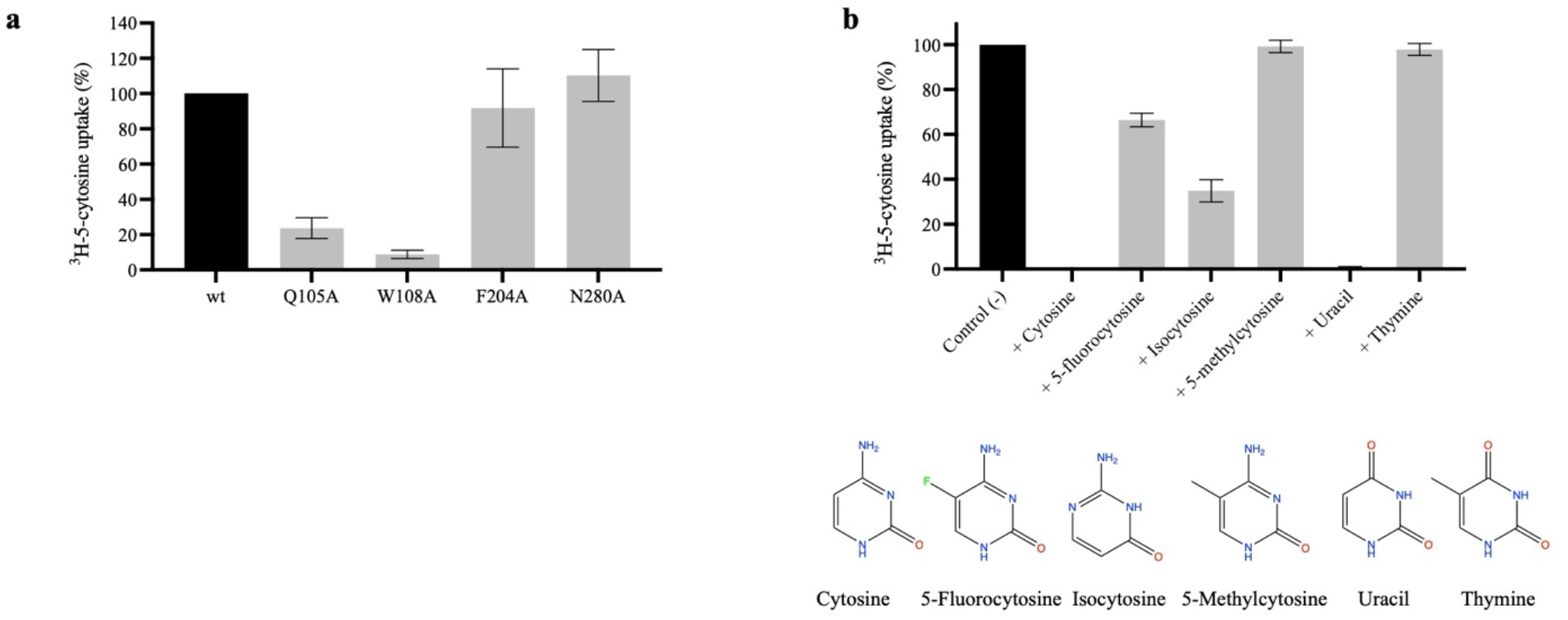
Functional characterisation of CodB. **a)** Uptake of ^3^H-5-cytosine by CodB mutants relative to the wild type protein. Uptake of ^3^H-5-cytosine was measured after 1 minute. Uptake for the wild type protein was set at 100%, and the mutants are shown as a percentage of this with error bars as s.e.m. of at least 4 experiments, each from a different culture. **b)** Inhibition of ^3^H-5-cytosine uptake in the presence of 0.1 mM of each respective inhibitor. Uptake of ^3^H-5-cytosine was measured after 1 minute with 0.1 mM inhibitor. Control (-) is uptake of ^3^H-5-cytosine with no inhibitor, normalised to 100%, results are visualised as % of control (-) with error bars as s.e.m. of triplicate experiments, each from a different culture. The chemical structures of the ligands are shown below the graph.

### Further interactions between bundle and hash domains

By investigating the pattern of conservation amongst CodB homologues we discovered that Arg216 on TM6b of the bundle domain and Tyr285 of TM8 of the hash domain are two of the most conserved residues (Supplementary Figure 5). Remarkably, these residues are within hydrogen bonding distance of one another at the cytoplasmic side of the protein. The high conservation suggests this interaction may be important for function. The same interaction is not found in Mhp1, however, in Mhp1 the arginine is replaced with a lysine (Lys232) and though the tyrosine is not conserved, the hydroxyl oxygen of Tyr324 one helix turn down is positioned such that a similar interaction would be possible (Supplementary Figure 6).

## Discussion

The structure of CodB we have elucidated here shows how the cytosine substrate makes specific hydrogen bonding interactions with the exposed main chain atoms at the breakpoint of TM6. Comparison with Mhp1 clearly shows that this is the common recognition site between these distantly related members of the NCS1 family. In other sodium-coupled members of the LeuT superfamily (Coleman *et al*., 2016; Faham *et al*., 2008; Malinauskaite *et al*., 2014; Penmatsa *et al*, 2015; Perez *et al*., 2012; Wahlgren *et al*., 2018; Yamashita *et al*., 2005) interactions with TM1 appear to be more important in anchoring the substrate than those of TM6 and those with TM6 to be more modulatory. In the NSS protein MhsT, for instance, the flexibility of the residues at the breakpoint of TM6 allow the accommodation of different amino acids (Focht *et al*, 2021). The sugar substrate in vSGLT is possibly the exception in not interacting with the main-chain of TM1 (Faham *et al*., 2008), but this is an inward-facing structure where the binding site is not fully formed. Interactions with the main-chain of TM1 are seen in the outward-facing sialic acid transporter, SiaT from the same family (Wahlgren *et al*., 2018). In CodB the only interaction between the substrate and TM1 is a stacking interaction with Phe33. Mutagenesis of the equivalent residue in Mhp1 led to the conclusion that the only function of this residue would be to shape the pocket (Simmons *et al*., 2014) and the position of Phe33 in CodB would support this conclusion.

Clearly the pi-stacking arrangement of the nucleobase between the two aromatic residues is also important. Interestingly, the mutation of Phe204 to Ala was much less drastic compared to the similar mutation of Trp108. A similar observation was made with an equivalent mutation in Mhp1 (Simmons *et al*., 2014). It seems likely that, while important for shaping the pocket, the prime binding site on the bundle domain involves the main chain atoms of TM6. Subtle changes caused by the interaction with the residues on the hash domain may then be important in allowing transport to occur. 5-Fluorocytosine could easily be accommodated with the same binding mode.

For alternating access to occur there are several changes that have been shown to take place in Mhp1. Following substrate binding, TM10 folds into the binding site, rotating around a conserved proline (Weyand *et al*., 2008). In our structure TM10 adopts a more open conformation. Although TM10 in CodB is one residue shorter than in Mhp1, given that the temperature factors are high for TM10 (Supplementary Figure 7) and the helix retains the proline on TM10 around which TM10 swivels (Figure 3) it seems likely that the lipid-like molecules that we observe in the density are preventing the conformation adopted in the substrate-bound form of Mhp1 rather than a substantial difference in mechanism. Molecular dynamics (Shimamura *et al*., 2010) and DEER (Kazmier *et al*., 2014b) both suggest this helix is very mobile in the outward-facing structure of Mhp1.

**Fig 7:**
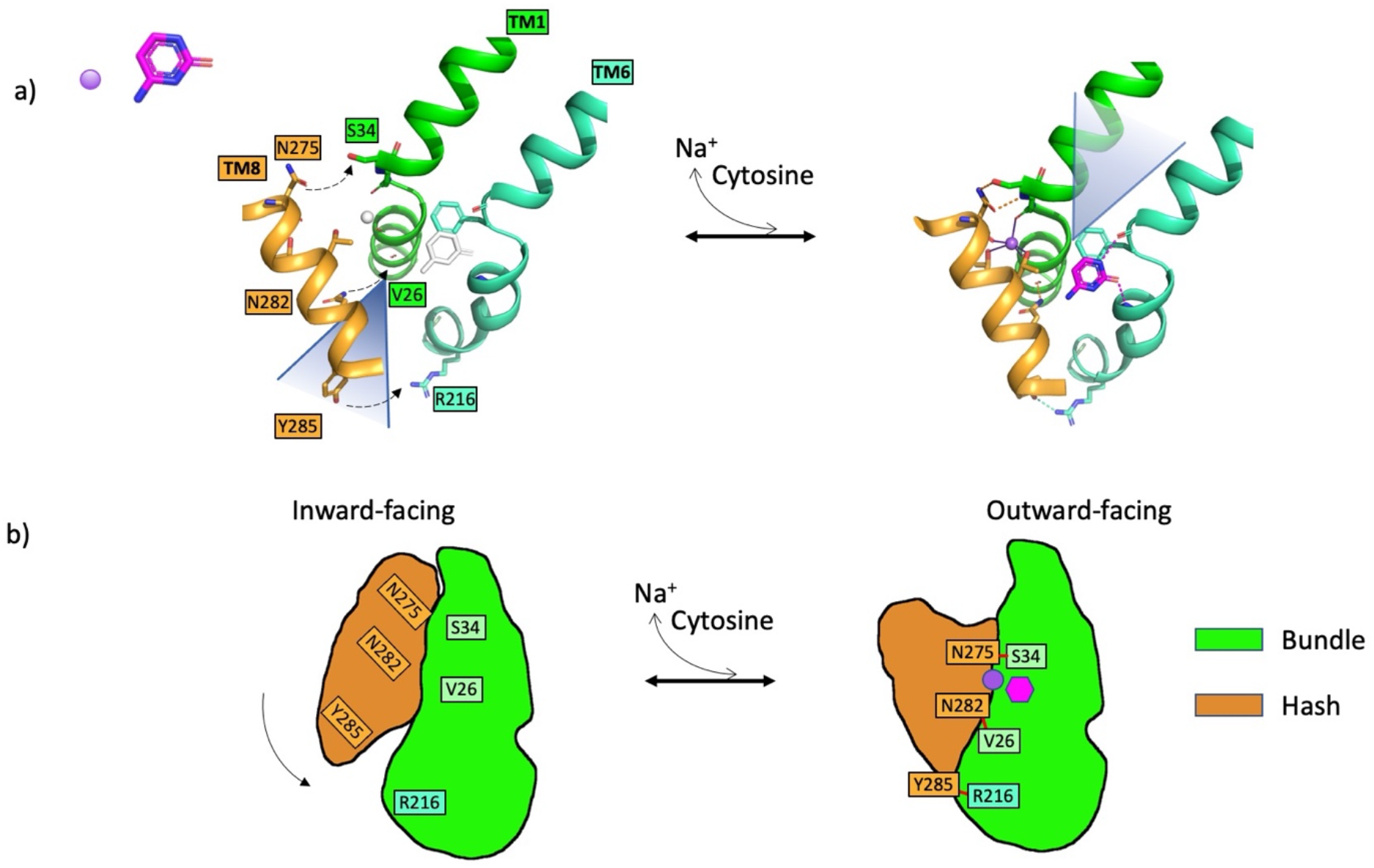
Mechanism of transition from inward to outward facing conformations in CodB. **a)** The putative movement of TM8 relative to the bundle as the protein transitions from the inward-facing state (left) to the outward-facing state (right). The position of TM8 in the inward-facing state has been modelled on the equivalent helix of Mhp1 in the inward-state (PDB code 2×79). In transitioning between the two states TM8 rotates around an axis approximately coincidental with TM3 bringing it much closer to TM1 so that the hydrogen bonding interactions seen in the outward-facing state (dashed lines) can form and the sodium ion can bind. These distances are too large for hydrogen bonds in the modelled inward-facing state. Cytosine will bind guided by residues on TM6 as well as TM3 (not shown) enabling conformational changes that will result in the transition back to the inward-facing state. The inward and outward facing clefts, which lie to the back of TM8 in the inward-facing state and to the front of TM1 in the outward-facing state are denoted by triangles behind and in front of the cartoons, respectively. **b)** A schematic showing the interactions between TM1 and TM8 acting like a zipper on the protein.

The second major conformational change in the transport cycle involves a rotation of the hash domain relative to the bundle domain (Shimamura *et al*., 2010). The rotation in Mhp1 is around an axis that is approximately coincidental with TM3 so that the movement of TM8 as the protein transitions from outward to inward facing is much greater than that of TM3. Intuitively, it would be thought that the sodium ion, which spans TM1 and TM8 in the outward-facing structure is likely to be important in shifting the equilibrium towards the outward-facing state. Certainly, the structure of Mhp1 in the presence of sodium ions, but without substrate is in the outward-facing state. However, DEER and mass spectrometry based studies of Mhp1 in detergent solution would appear to go against this, (Calabrese *et al*, 2017; Kazmier *et al*., 2014a) with both indicating that the presence of the substrate as well as sodium ions are required to drive a conformational change from inward to outward-facing states in contrast to a similar study carried out for LeuT (Kazmier *et al*., 2014b). It may well be that these results are affected by these experiments being carried out in a non-lipid environment in the absence of a membrane potential as was suggested for similar experiments on vSGLT (Paz *et al*, 2018). It certainly seems likely that subtle changes in the energetics of the system are likely to influence the conformational state of the protein. In Mhp1, Asn318 on TM8 makes an important bidentate hydrogen-bonding interaction with the substrate so this is likely to influence the conformational change. In CodB, there is only a water-mediated interaction between TM8 and the cytosine. Instead, there are direct hydrogen bonding interactions, between TM8 and TM1, one involving Asn275, which is also a ligand to the sodium ion and the other from Asn282 which is just below the sodium ion. These residues may affect the activity of the protein, albeit subtly, by making it energetically more favourable to adopt the outward-facing state in the presence of sodium ions. It is noteworthy that in the sialic acid transporter, SiaT, the equivalent residue to Asn282 is involved in a second sodium-binding site, which appears to modulate activity (Wahlgren *et al*., 2018) and in MhsT the equivalent residue is a serine, which is also involved in a hydrogen bonding interaction with a serine on TM1 (Malinauskaite *et al*., 2014). In general, the hydrogen bonding arrangement between residues of the hash domain and residues of the bundle domain in CodB differ widely from those in Mhp1. In Mhp1 there are no residues from TM8 that are involved in direct hydrogen bonding interactions with the bundle domain but two residues on TM3. The first is Gln121, which although it hydrogen bonds to the substrate is also within hydrogen bonding distance of Gln42 in the substrate-bound structure. The second is Lys110 (Supplementary Figure 6), which has been suggested to mimic a sodium ion binding site that is observed in the betaine transporter, BetP (Khafizov *et al*, 2012). The lysine amino group interacts with both the hydroxyl oxygen of Ser226 of TM6 of the bundle domain and Ser114 on TM3 of the hash domain. In CodB, the lysine is replaced with a hydrophobic residue, so cannot form the same interaction. Interestingly, the equivalent of Ser114 with which the lysine interacts in Mhp1 is Gln105, which forms hydrogen bonds with the ligand.

Given the conservation of Tyr285 and Arg216 amongst putative CodB homologues from different organisms, the interaction between them appears to be important. This interaction is reminiscent of that between Tyr268 and Gln361 of LeuT (Yamashita *et al*., 2005), which is conserved in the NSS family. The mutation of Tyr268 in LeuT favours the inward-open structure (Kazmier *et al*., 2014b; Krishnamurthy & Gouaux, 2012). CodB also resembles LeuT in that Arg216 also interacts with the N-terminus (Supplementary Figure 6). Though there is no conservation in the residues involved, the interaction between Arg5 at the N-terminus of LeuT and Asp369 within the scaffold domain is important in the mechanism of LeuT and NSS transporters (Kniazeff *et al*, 2008; Krishnamurthy & Gouaux, 2012). It has been shown for other NCS1 members that the N-terminus affects the mechanism and specificity of the transporters (Papadaki *et al*, 2019). It therefore seems that while each of the proteins has important interactions linking the two domains, the exact mode widely varies amongst them.

In conclusion, the high-resolution structure of CodB with cytosine in combination with site-directed mutagenesis has enabled us to understand substrate binding in CodB and see that 5-fluorocytosine could easily be accommodated in the binding site. It also illustrates the importance of the interaction between the substrate and TM6 in the NCS1 family. The structural analysis highlights how the interaction with the sodium ion and substrate are separated, with the sodium ion binding to TM1 and the substrate primarily interacting with TM6 (Fig 7), unlike the arrangement in other characterised members of the superfamily. Whether this can be correlated with the larger movements of TM1 seen in other members of the superfamily during the transport cycle remains to be seen. It seems likely that the three hydrogen bonding interactions between residues of TM1 and TM8 discussed above, will also influence the mechanism. Presumably these interactions will stabilise the outward-facing state of the protein in readiness for the cytosine to bind (Fig 7). The structural analysis provides further insight into how a common mechanism of sodium-coupled symport in this superfamily is modulated by structurally similar proteins in diverse ways.

## Materials and Methods

### Expression and Protein Purification

The gene encoding for the cytosine permease CodB from *P. vulgaris*, codon optimised for expression in *E coli* was purchased as a gBlock (Integrated DNA Technologies). This was inserted into a modified version of the expression vector, pWaldo GFPd (Drew *et al*., 2006) in which the TEV protease site had been altered to a site for recognition by 3C protease. Site-directed mutations were introduced by PCR (Quikchange, Agilent Technologies). CodB-GFP fusions were expressed in *E. coli* Lemo21 (DE3) cells following the MemStar procedure (Lee *et al*, 2014). Briefly, cultures were grown at 37 °C, 200 rpm, in PASM-5052 media supplemented with 0.1 mM rhamnose. When cultures reached OD_600_=0.5, the temperature was dropped to 25 °C and 0.4 mM IPTG was added for protein induction overnight.

Cell pellets were harvested by centrifugation and resuspended in PBS (137 mM NaCl, 2.7 mM KCl, 10 mM Na_2_HPO_4_, 1.8 mM KH_2_PO_4_) with 1 mM MgCl_2_, DNaseI, and 0.5 mM 4-benzenesulfonyl fluoride hydrochloride (AEBSF) and disrupted by passing three times through a cell disruptor at 25 kPsi. Cell lysate was centrifuged at 24,000 g at 4 °C for 12 minutes to remove insoluble cell debris, and the supernatant was subjected to ultracentrifugation at 200,000 g, 4 °C for 45 minutes. Membrane pellets were resuspended in PBS, 15 mL per 1 l of culture, snap frozen in liquid nitrogen, and then stored -80 °C.

For crystallisation membranes from 3 L of culture were solubilised in 1x PBS, 150 mM NaCl, 1% DDM for 2h at 4°C and ultracentrifuged for 45 mins, 4 °C, 200,000g to remove insoluble material. Imidazole was added to 20 mM, and the membrane suspension was mixed with 1 ml of Ni-NTA Superflow resin (Qiagen) per 1mg of GFP–His8 and incubated for 3 hours at 4 °C. Slurry was decanted into a glass Econo-Column (Bio-Rad) and washed with 5 Column Volumes (CV) of 1x PBS, 150 mM NaCl, 20 mM imidazole, 0.1% DDM, then 5 CV of 20 mM TRIS pH 7.5, 150 mM NaCl, 30 mM imidazole, 0.03% DDM, 1mM cytosine. Protein was left on the column overnight with a 1:1 stoichiometry of 3C protease at 4 °C. Cleaved protein was eluted into fractions corresponding to 1CV and passed over a 5 ml HisTrap equilibrated with 20 mM TRIS pH 7.5, 150 mM NaCl, 30 mM imidazole, 0.03% DDM to remove contaminants. Protein was concentrated to 32 mg/ml using centrifugal concentrators (Sartorius) with a relative molecular mass cut-off of 100K.

### Transport Time Course

CodB was expressed in Lemo21(DE3) cells as above with 25ml culture volumes. Following centrifugation of the cultures at 2,600 g for 10 minutes at 20 °C, the supernatant was removed and the pellet was resuspended in 5 ml 5 mM MES pH 6.6, 150 mM KCl. This was repeated three times. Cells were resuspended to give a final concentration of OD_600_ of 2 in 1200 μL of either 5 mM MES pH 6.6, 150 mM NaCl or 5 mM MES pH 6.6, 150 mM choline chloride. 6 µl of 6.25 µM ^3^H-5-cytosine (20 Ci mmol^-1^; American Radiolabelled Chemicals) was added to samples and the cells were incubated at 37 °C with shaking at 900rpm for times of 30 seconds, 1 minute, 2 minutes, 5, minutes, 10 minutes, or 20 minutes. At the stated timepoint, 200 μL of cells were centrifuged at 13,000rpm for 30 seconds at 20 °C, the supernatant was removed and the pellet was resuspended in 200 µl stop buffer (5 mM MES pH 6.6, 150 mM KCl, 1 mM cytosine) and added to a 0.2 µm Whatman cellulose nitrate membrane filter under vacuum followed by immediate washing with 4 × 2 mL 0.1 M LiCl. Each filter was placed in 10 ml Emulsifier Safe scintillation fluid and counted using a Tri-CarbA4810TR Liquid Scintillation Analyzer (Perkin Elmer). CodB concentration was quantified based on the GFP fluorescence. Lemo21 cells with no CodB overexpression were used as a background, with 1 µl of 6.25 µM ^3^H-5-cytosine used to calibrate counts. Experiments were performed in triplicate with fresh cultures.

### Inhibition assay

Cells were prepared as described previously and resuspended to an OD_600_ of 2 in 200 μL to give a final concentration with 5 mM MES pH 6.6, 150 mM NaCl, 0.1 mM potential inhibitor. 1 μL of 6.25 μM ^3^H-5-cytosine was added and the mixture incubated at 37 °C with shaking at 900 rpm for 1 minute before centrifuging at 16,000 g for 1 minute at 20 °C. The supernatant was removed and the pellet was resuspended in 200 μL stop buffer (5 mM MES pH 6.6, 150 mM KCl, 1 mM cytosine) and added to a 0.2 μm Whatman cellulose nitrate membrane filter under vacuum followed by immediate washing with 4 × 2 mL 0.1 M LiCl. All filters were dissolved in 10 ml Emulsifier Safe scintillation fluid and counted using a Tri-Carb A4810TR Liquid Scintillation Analyzer (Perkin Elmer). Experiments were performed in triplicate with fresh cultures.

### GFP-TS

The GFP-TS assay was carried out following the published protocol (Nji *et al*., 2018). 150 µl of *E. coli* membrane with overexpressed CodB was diluted 1:10 in 20 mM TRIS pH 7.5, 150 mM NaCl, 1% DDM, 1% octyl-β-D-glucoside, (β-OG), 1 mM of the molecule to be tested and left mixing at 4 °C for 1 h then aliquoted into 150 µl fractions. Aliquots were subjected to various temperatures, 4 °C, 20 °C, 25 °C, 30 °C, 35 °C, 40 °C, 45 °C, 50 °C, 60 °C for 10 minutes then spun at 16,000 g for 30 minutes. 100 µl of supernatant was transferred to a 96 well black-walled plate and GFP measurements were taken. The apparent T_m_ for each titration was calculated by plotting the normalised average GFP fluorescence intensity from two technical repeats at each temperature and fitting the curves to a sigmoidal dose–response equation (variable slope) by GraphPad Prism software (version 9.0). Values reported are the averaged mean ± s.e.m. of the fit from n = 2 independent titrations.

To generate an approximate K_d_ 150 µL of *E. coli* membrane was solubilised as before but cytosine was added at a final concentration between 0-1000 µM. Aliquots were put at 35 °C for 10 minutes and spun at 16,000 g for 30 minutes. 100 µl of supernatant was transferred to a 96 well black plate and GFP measurements were taken. Binding curve was fitted by nonlinear regression (one site, total binding) by GraphPad Prism software (version 9.0), and the values reported are the averaged mean ± s.e.m. of the fit from n = 3 independent titrations.

### Crystallisation and Structural Determination

Protein at a concentration of 32mg/ml was subjected to crystallisation using the lipidic cubic phase method of crystallisation (Caffrey & Cherezov, 2009). The CodB protein with 1mM cytosine was mixed with monoolein at 60:40 (w/w) ratio using a coupled syringe device (Molecular Dimensions Ltd UK) and crystallisation trials were set up at 20°C using glass sandwich plates using a Mosquito Robot. Crystals appeared in condition G5 of MemGoldMeso (Molecular Dimensions Ltd) in glass sandwich plates, which contained 0.1 M sodium cacodylate pH 6.5, 0.45 M NaCl, 39% PEG400. Crystals were cryo-cooled in liquid nitrogen.

X-ray diffraction data were collected at beamline I24 at Diamond Light Source, UK. Initially a data set was collected that was processed at 3.6Å resolution but subsequently a higher resolution data set was collected. Data were processed using DIALS (Waterman *et al*, 2016) through the Xia2 pipeline (Winter *et al*, 2013). Processed data was then scaled and merged in AIMLESS (Evans & Murshudov, 2013) in the CCP4 suite (Collaborative Computational Project Number 4, 1994). The resolution cut-off was chosen based on where the CC_0.5_ fell below 0.5. The structure was solved from the 3.6Å resolution data set using MR_ROSETTA (DiMaio *et al*, 2011) in the PHENIX package (Liebschner *et al*, 2019) basing the search on the outward-facing structure of Mhp1 (2JLN; (Weyand *et al*., 2008)). Refinement was carried out with PHENIX.REFINE (Afonine *et al*, 2012) interspersed with model building in Coot (Emsley & Cowtan, 2004) initially against the low resolution data set but subsequently against the high resolution data set.

Superpositions were performed in Chimera (Pettersen *et al*, 2004) maintaining the default cut-off of 2Å for pruning matching C_α_ atoms and structural images were prepared in PyMol (Delano, 2002). Images involving electron density were prepared in CCP4mg (McNicholas *et al*, 2011).

To obtain the sequence alignment for proteins similar to CodB from P. vulgaris a BLAST search (Altschul *et al*, 1990) was carried out at the EBI (Madeira *et al*, 2019) against the Uniref90 database from the Uniref clusters (Suzek *et al*, 2015) selecting 200 sequences. These were aligned using Clustal Omega (Sievers *et al*, 2011) and imported into Jalview (Waterhouse *et al*, 2009). Similar sequences were removed using the “Remove Redundancy” tool in Jalview. The sequence alignment figure was based on the image output from Jalview.

## Acknowledgments

We thank the staff at beamline I24 at DLS and the technical support in the School of Life Sciences, University of Warwick. We thank Dr Phill Stansfeld for commenting on the manuscript. Funding: The project was supported by a BBSRC MIBTP studentship to CEH. DHB was supported by MRC (MR/P010393/1).

## Author contributions

The project was initiated by AC. CEH carried out the experiments, with help and supervision by AC and DHB. MS contributed to the GFP-TS assays. The paper was written by AC with contributions from CEH and DHB.

## Data and materials availability

The structure and data have been deposited in the RSCB with accession number xxx.

## Supplementary Figures

**Fig S1:**
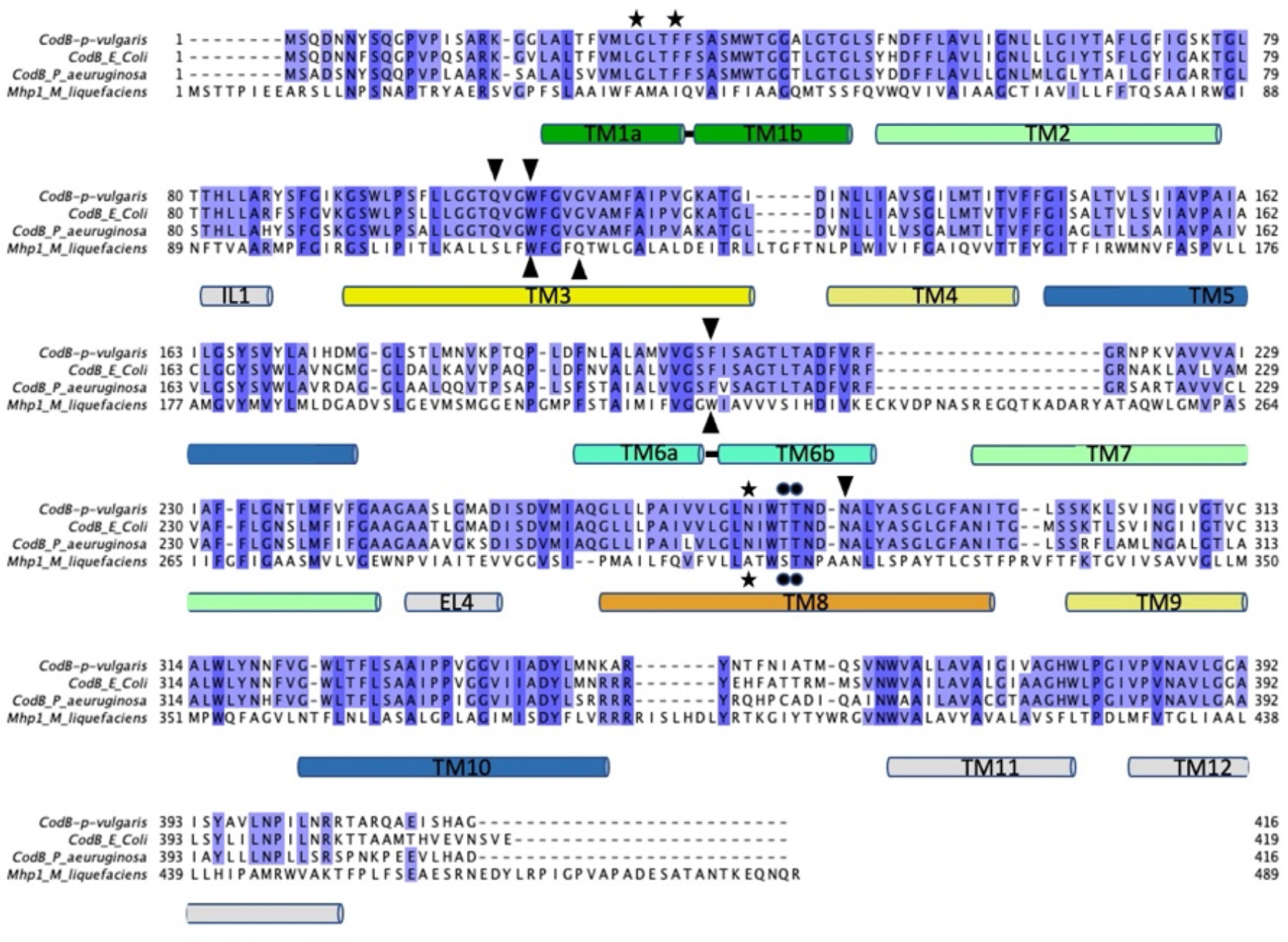
Sequence alignment of CodB and Mhp1. Alignment of sequences of CodB from *P. vulgaris, E. coli* and *P. aeruginosa* with Mhp1 from *M. liquefaciens* shaded according to sequence conservation. The sequences were aligned with Muscle (Edgar, 2004) with manual adjustments based on the respective structures. The secondary structure is shown for CodB. Residues interacting with the substrate are denoted by triangles (▾ for CodB and ▴ for Mhp1), residues interacting with the sodium ion through the carbonyl oxygen are shown as ★ and those interacting through the side chain are denoted by •.

**Fig S2:**
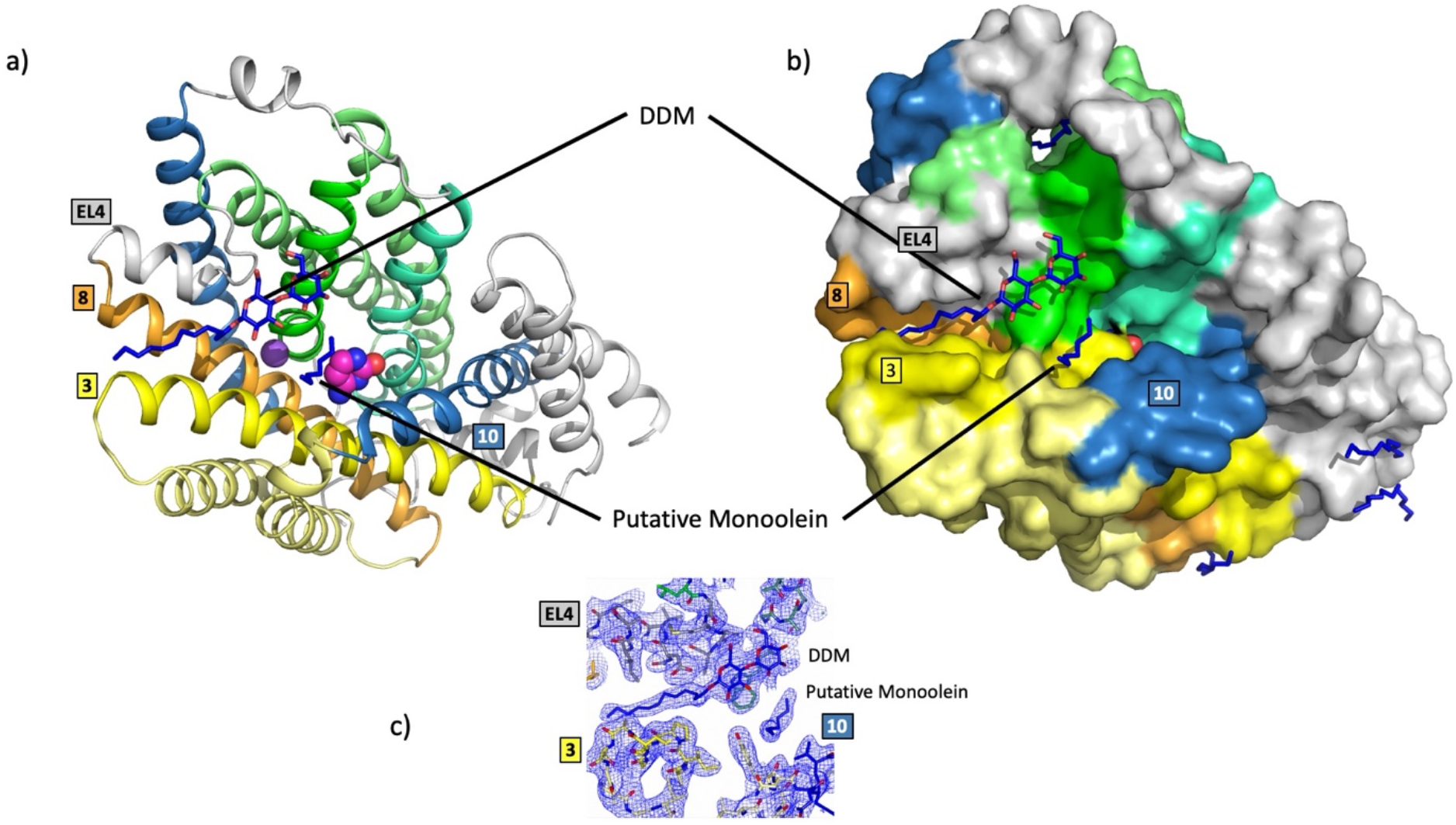
Detergent and lipid binding in CodB. As Fig 2b but showing positions of monoolein and DDM that have been putatively modelled in the binding site. Lipids are shown with blue carbon atoms. **a)** cartoon representation, **b)** surface representation. **c)** Electron density for lipid like molecules in the cavity. The 2mFo-DFc map is based on phases from the refined structure and contoured at 0.75σ.

**Fig S3:**
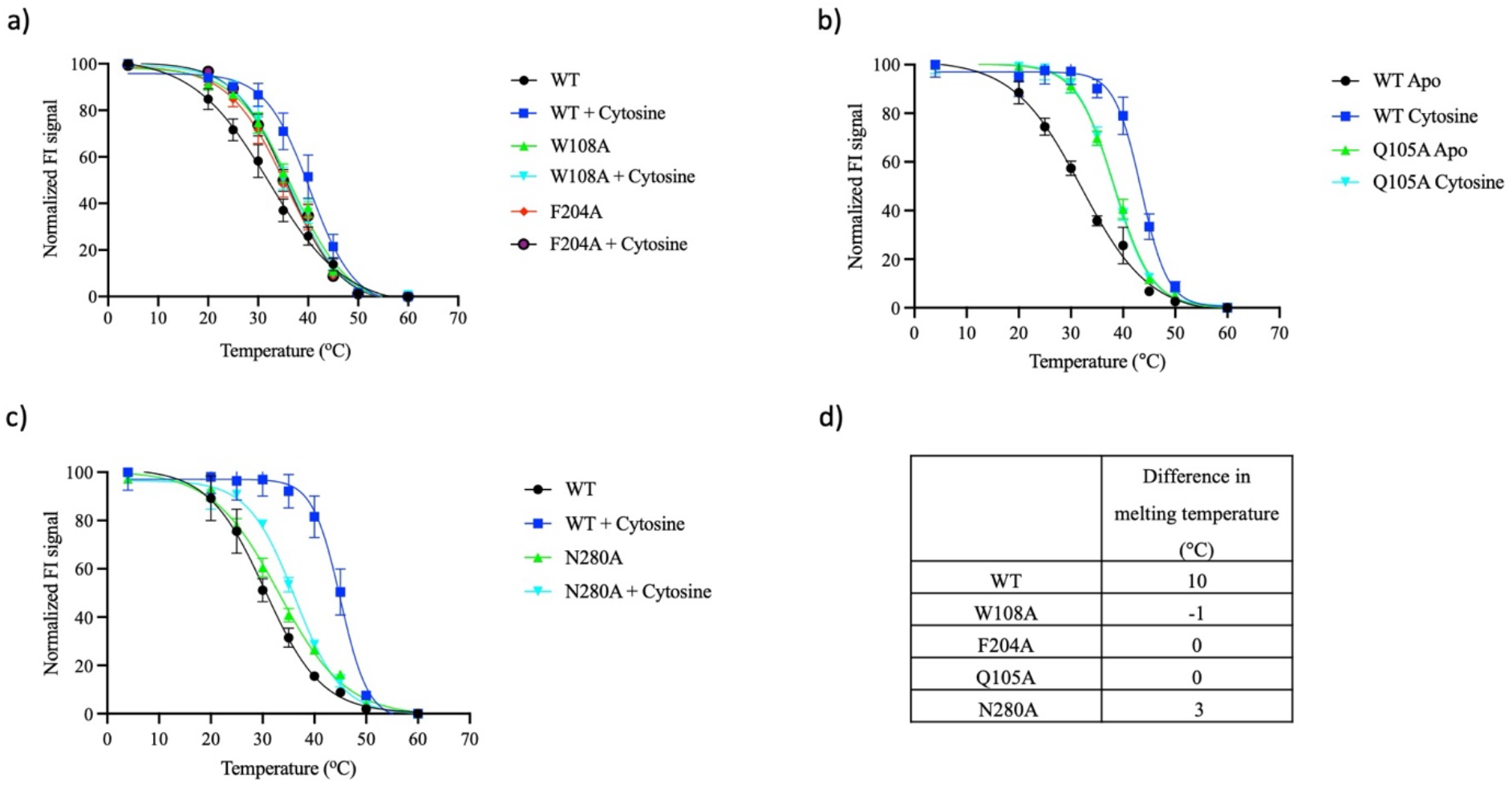
Characterisation of CodB mutants using the thermostability assay. **a**,**b**,**c)** Melting curves for wild type and mutant proteins with and without cytosine bound using the GFP-TS assay. Values reported are the averaged mean ± s.e.m. of the fit from n = 2 independent titrations. **d)** Table showing differences in melting temperatures between protein with and without cytosine bound.

**Fig S4:**
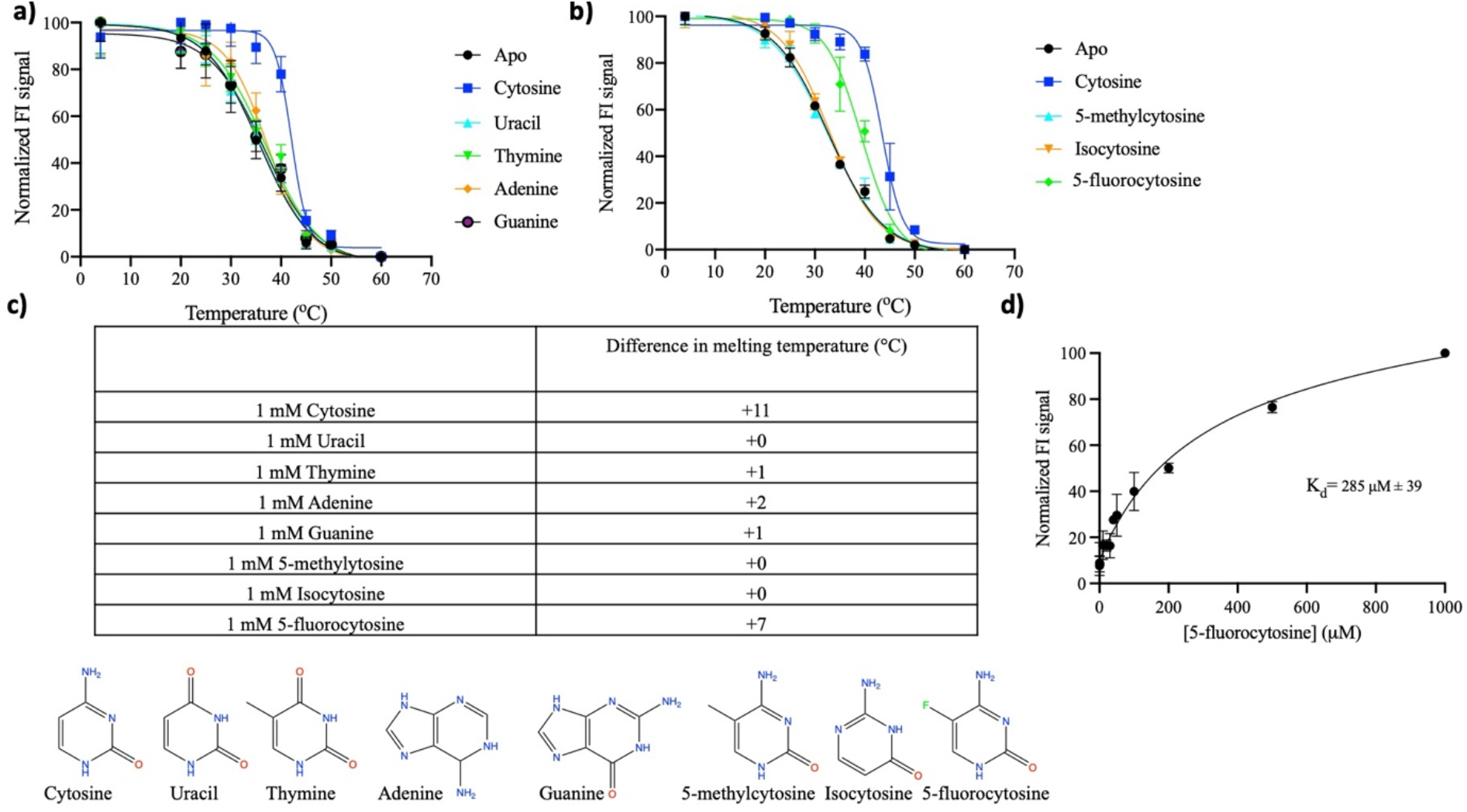
Characterisation of nucleobase binding to CodB using the thermostability assay. **a**,**b)** Melting curves for the wild type CodB in the presence and absence of selected nucleobases using the GFP-TS assay. Values reported are the averaged mean ± s.e.m. of the fit from n = 2 independent titrations. **c)** Table showing differences in melting temperatures upon addition of the specified nucleobase. d) Binding affinity of CodB for 5-fluorocytosine as measured using the thermostability assay. The K_d_ was estimated to be 285 ± 39 µM. The measurements are the average of 3 independent titrations with error bars of the s.e.m.

**Fig S5:**
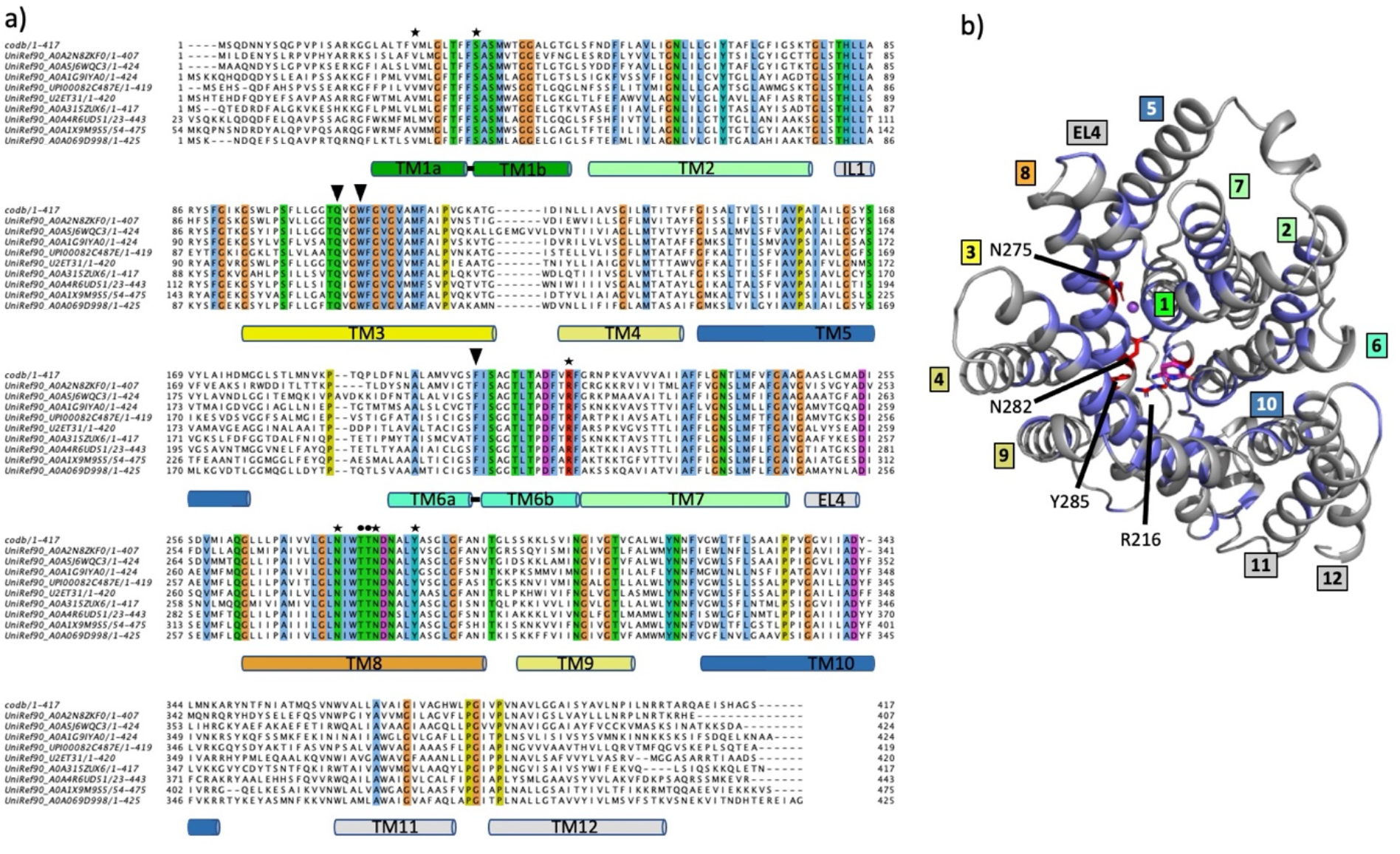
Sequence conservation in CodB. **a)** A BLAST search was carried out against UniProt clusters (UniRef90). The most diverse sequences from the 200 sequences deriving from that search (see methods) are shown. Residues that are identical in all 10 sequences are highlighted coloured according to ClustalX. Residues interacting with the ligand are shown by a solid triangle (▾), those interacting with the sodium ion through their side chains are shown as a • and those involved in hydrogen bonds between the helices of the bundle and hash domain respectively are denoted with a ✶. As more diverse sequences are added R216 and Y285 remain constant while there is more divergence for other residues. **b)** Conservation plotted on the structure of CodB. Residues that are identical in (a) have been coloured blue with N275, N282, R216 and Y285 depicted as red sticks.

**Fig S6:**
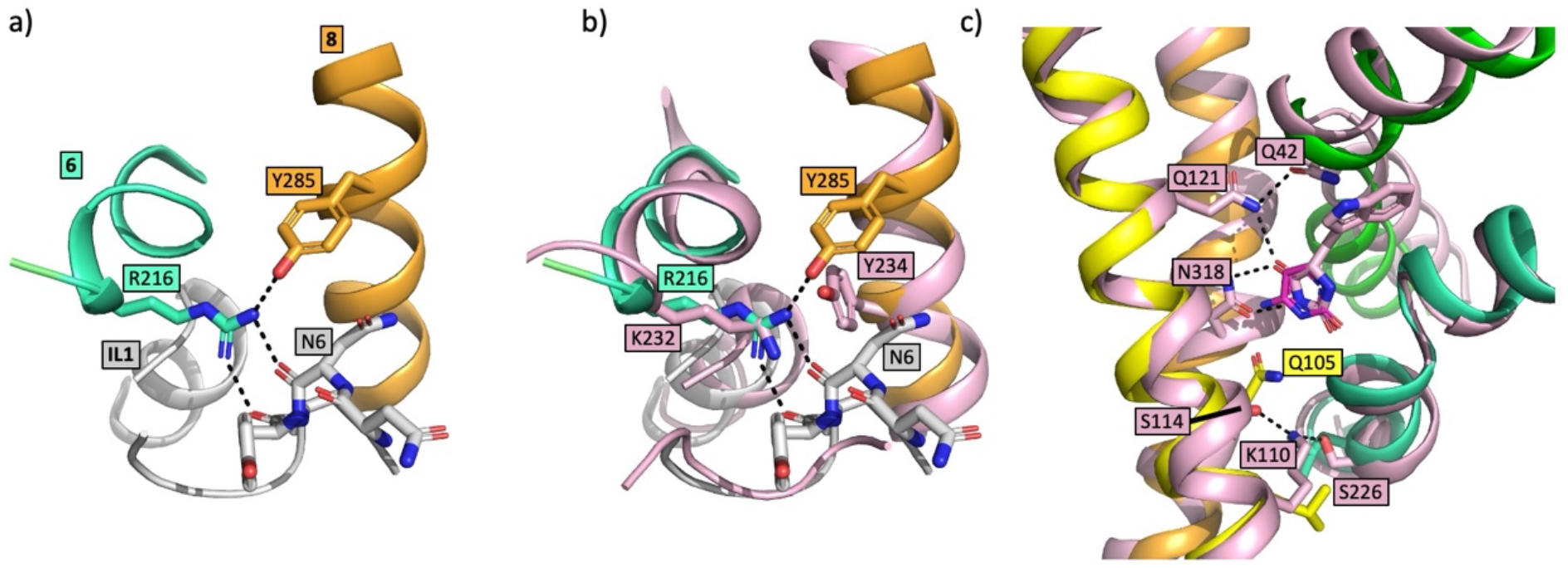
Interaction between residues of the bundle and hash domains in CodB and Mhp1. **a)** CodB: Interaction between Arg216 and Tyr285 coloured as in Figure 2. Hydrogen bonds are shown as dashed lines. **b)** As (a) with Mhp1 superposed on the structure. In Mhp1 Arg216 is replaced by a lysine and while Tyr285 is not conserved, the hydroxyl group of another tyrosine could interact with the lysine. **c)** Comparison between the hydrogen bonding arrangement in CodB and Mhp1 with potential hydrogen bonds shown for Mhp1

**Fig S7:**
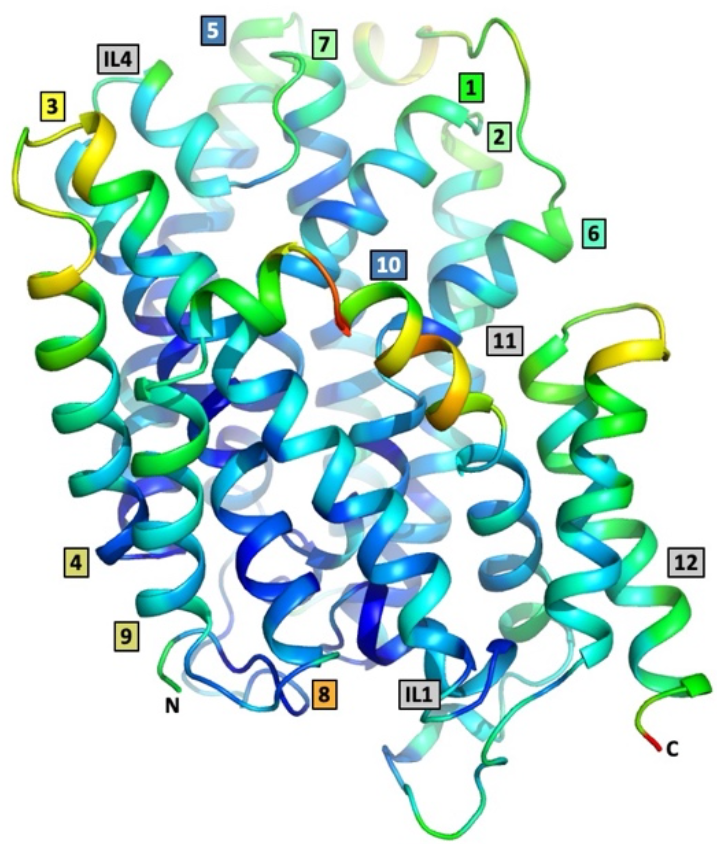
The structure of CodB coloured by temperature factors. The higher the temperature factor the warmer the colour.

